# Human 5′ UTR design and variant effect prediction from a massively parallel translation assay

**DOI:** 10.1101/310375

**Authors:** Paul J. Sample, Ban Wang, David W. Reid, Vlad Presnyak, Iain McFadyen, David R. Morris, Georg Seelig

## Abstract

Predicting the impact of *cis*-regulatory sequence on gene expression is a foundational challenge for biology. We combine polysome profiling of hundreds of thousands of randomized 5′ UTRs with deep learning to build a predictive model that relates human 5′ UTR sequence to translation. Together with a genetic algorithm, we use the model to engineer new 5′ UTRs that accurately target specified levels of ribosome loading, providing the ability to tune sequences for optimal protein expression. We show that the same approach can be extended to chemically modified RNA, an important feature for applications in mRNA therapeutics and synthetic biology. We test 35,000 truncated human 5′ UTRs and 3,577 naturally-occurring variants and show that the model accurately predicts ribosome loading of these sequences. Finally, we provide evidence of 47 SNVs associated with human diseases that cause a significant change in ribosome loading and thus a plausible molecular basis for disease.

---

The sequence of the 5′ untranslated region (5′ UTR) is a primary determinant of translation efficiency (*1, 2*). While many cis-regulatory elements within human 5′ UTRs have been characterized individually, the field still lacks a means to accurately predict protein expression from 5′ UTR sequence alone, limiting the ability to estimate the effects of genome-encoded variants and the ability to engineer 5′ UTRs for precise translation control. Massively parallel reporter assays (MPRAs) – methods that assess thousands to millions of sequence variants in a single experiment – coupled with machine learning have proven useful in addressing similar voids by producing quantitative biological insight that would be difficult to achieve through traditional approaches (*3–9*).

We report the development of an MPRA that measures the translation of hundreds of thousands of randomized 5′ UTRs via polysome profiling and RNA sequencing. We then use the data to train a convolutional neural network (CNN) that can predict ribosome loading from sequence alone. Earlier MPRAs designed to learn aspects of 5′ UTR *cis*-regulation relied on FACS (*10, 11*) or growth selection (*12*) to stratify libraries by activity. These techniques require the expression of a single library variant per cell that must be transcribed within the cell from a DNA template, making it difficult to distinguish between the effects of transcriptional and translational control. Polysome profiling (*13*) overcomes this limitation by enabling single cells to translate tens to hundreds of in *vitro* transcribed (IVT) and transfected mRNA variants. Polysome profiling has been used extensively to measure translation of native RNA isoforms (*14, 15*) but isolating the role of 5′ UTR regulation is difficult due to differences in the size and sequence of the concomitant coding sequences and 3′ UTRs. To build a model capable of predicting the ribosome loading of human 5′ UTR variants and designing new 5′ UTRs for targeted expression (**Fig. 1A**), we first created a 300,000-member gene library with random 5′ UTRs but constant eGFP coding sequence and 3′ UTR (**Fig. 1B**). Specifically, the 5′ UTR of each construct begins with 25 nucleotides of defined sequence used for PCR amplification, followed by 50 nucleotides of fully random sequence before the eGFP coding sequence. HEK293T cells were transfected with IVT library mRNA and harvested after 12 hours. Polysome fractions were collected and sequenced (**Fig. S1A**). For a given UTR, the relative counts per fraction were multiplied by the number of ribosomes associated with each fraction and then summed to obtain a measured Mean Ribosome Load (MRL). We focused on the first 50 bases upstream of the CDS to specifically investigate the regulatory signals that mediate the initiation of translation beyond ribosomal recruitment to the 5′ cap.

**Figure 1.**
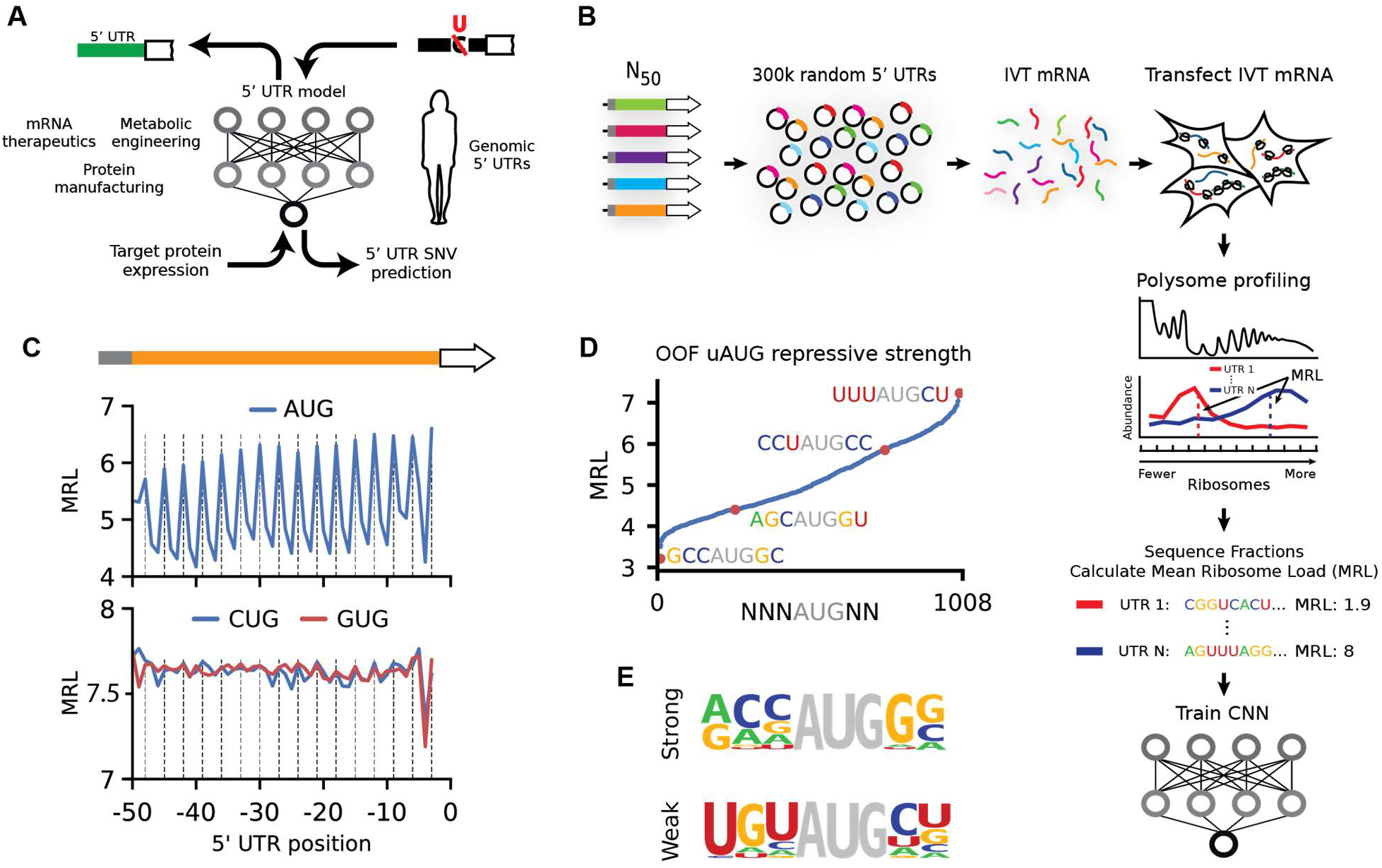
Experiment overview. **(A)** A 5′ UTR model capable of predicting translation from sequence is used to evaluate the effect of 5′ UTR SNVs and to engineer new sequences for optimal protein expression. **(B)** A library of 300,000 random 50-mers serve as 5′ UTRs for eGFP. Cells transfected with library IVT mRNA were grown for 12 hours before polysome profiling. Read counts per fraction were used to calculate Mean Ribosome Loads (MRL) for each UTR and the resulting data were used to train a convolutional neural network (CNN). **(C)** Out-of-frame upstream AUGs (uAUGs) reduce ribosome loading (vertical lines indicate positions that are in-frame with the eGFP CDS). A similar but much weaker periodicity is observed for CUG and GUG. **(D)** The repressive strength of all out-of-frame variations of NNNATGNN. **(E)** Nucleotide frequencies were calculated for the 20 most repressive (‘strong’) and least repressive (‘weak’) TIS sequences.

To validate our approach, we asked whether it captured known aspects of translation regulation. Translation initiation is largely dependent on start codons and their context and position relative to a CDS (*12, 16*). Our data clearly show the expected decrease in ribosome loading for sequences with either out-of-frame upstream start codons (uAUGs) (**Fig. 1C**) or upstream open reading frames (uORFs) (**Fig. S2B**) (*17, 18*). Interestingly, we observed only a minimal use of CUG and GUG as alternative start codons (**Fig. 1C, S3, and S4**) unlike other reports that show widespread usage of non-AUG start sites (*15, 19, 20*), possibly because these alternative start codons are used more often under stress conditions (*21*). The region surrounding the start codon, known as the translation initiation site (TIS) or the Kozak sequence, is a primary determinant of whether a ribosome will begin translation. We scored the repressive strength of all out-of-frame TISs by finding the mean MRL of sequences with all permutations of NNNAUGNN (except where NNN is AUG) (**Fig. 1D**). Using the 20 most repressive and 20 least repressive sequences, we calculated nucleotide frequencies for the strongest and weakest TISs. This analysis recapitulated the importance of a purine (A or G) at position -3 relative to AUG and a G at +4 (**Fig. 1E**) (*10, 22, 23*). Ultimately, these data suggest that each TIS sequence can uniquely tune translation initiation to a fine degree. Translation initiation and elongation is also affected by RNA secondary structure that forms within 5′ UTRs and coding sequences, with strong structures showing the most negative effect on translation (*16, 24*). By calculating UTR minimum free energies (MFE) (25) and comparing them to UTR MRLs, we captured and quantitated this repressive effect of secondary structure on ribosome load (**Fig. S2C**) (*16, 24*).

Next, we set out to develop a model that could quantitatively capture the relationship between 5′ UTR sequences and their associated MRLs. To this end, we trained a convolutional neural network (CNN) with 280,000 of the 300,000-member eGFP library. The remaining 20,000 sequences were withheld for testing. After an exhaustive grid search to find optimal hyperparameters (Online Methods) (**Fig. 2A**), the model could explain 93% of MRL variation in the test set (**Fig. 2B**). A model trained on data from a biological replicate performed similarly (**Fig. S5A**). By comparison, a position-specific 5-mer linear model could only explain 66% of the variation in the test set (**Fig. S6**).

**Figure 2.**
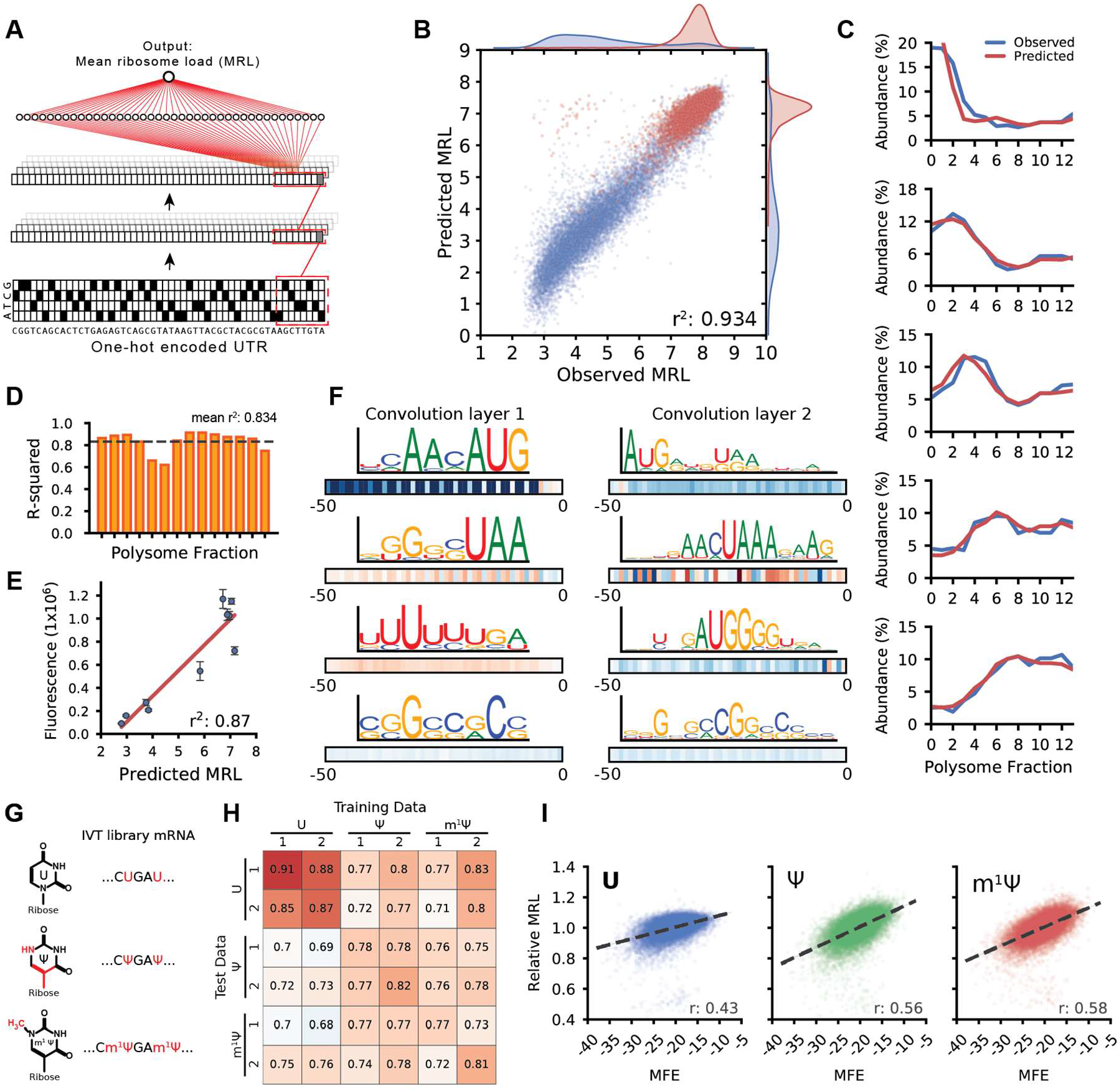
Modeling 5′ UTR sequences and ribosome loading. **(A)** A one-hot encoded 5′ UTR sequence is fed into a CNN composed of three convolution layers and a fully connected layer to produce a linear output predicting MRL. **(B)** A model trained on 280,000 UTRs and tested on 20,000 held-out sequences could explain 93% of the variability in observed MRLs. Blue dots represent sequences with an uAUG while red dots represent sequences without uAUG **(C)** A similar model was trained to predict the polysome profile distribution of an individual 5′ UTR. The observed (blue) and predicted (red) polysome distribution of 6 example UTRs spanning MRLs from 4 to 8 (top to bottom) are shown. **(D)** The performance of the polysome profile model per fraction ranged from an r^2^ of 0.621 to 0.915 and an average of 0.834 across all fractions. **(E)** eGFP expression for ten UTRs selected from the library were evaluated via eGFP fluorescence using IncuCyte live cell imaging. Predicted MRL and fluorescence are highly correlated (r^2^: 0.87). **(F)** Visualization of four out of 120 filters from the first convolution layer (left) and four out of 120 filters from the second convolution layer. Boxes below show correlation (Pearson r) between filter activation and MRL at each UTR position. Filters learned important regulatory motifs such as start and stop codons, uORFs, and GC-rich regions likely involved in secondary structure formation. **(G)** IVT mRNA from the eGFP library were generated with pseudouridine (Ψ) or 1-methylpseudouridine (m^1^ Ψ) in place of uridine (U) and evaluated by polysome profiling and modeling. **(H)** Model performance trained and tested on different data sets (r-squared). The unmodified RNA (U) models perform best with U data, while the Ψ and m^1^ Ψ models perform equally well with Ψ and m^1^ Ψ test data. **(I)** Ribosome loading as a function of MFE. U is less dependent on secondary structure than Ψ and m^1^ Ψ (Pearson r: 0.43, 0.56, and 0.58, respectively).

So far, we used MRL as a simple measure for translation but the raw data also captures how often a given sequence occurs in each polysome fraction. We thus set out to build a model capable of predicting the full polysome distribution for a given sequence. Using a similar network architecture but with 14 linear outputs representing the polysome fractions (**Fig. S7B**), the model captured the relationship between 5′ UTR sequence and distribution of ribosome occupancy on held out test data remarkably well (**Fig. 2C**), explaining an average of 83% of variation across all fractions (**Fig. 2D**). To test whether the mean ribosome load prediction corresponds to actual protein expression, we selected and synthesized mRNAs containing 10 different UTRs from the library with a wide range of observed MRLs. We then transfected these mRNAs into HEK293 cells and measured eGFP fluorescence using IncuCyte live cell imaging. Fluorescence and predicted MRL were highly correlated (r^2^: 0.87) and the most poorly translated sequence showed 15-fold less fluorescence than the best (**Fig. 2E**). Finally, to learn whether the model would generalize to other coding sequences, we built a separate degenerate 5′ UTR mRNA library with an mCherry CDS replacing eGFP. Following the polysome profiling and modeling procedure as above, we found that the model, although only trained on the eGFP library, still performed well, explaining 77% and 78% of the variation in MRL for two replicates of this new reporter library (**Fig. S5**). The decrease in accuracy is explained in part due to differences between the eGFP and mCherry polysome profiling protocols (Online Protocols).

To aid interpretation of the model we applied visualization techniques developed in computer vision and recently popularized in computational biology (*4, 8, 26*). Visualization of the filters in the first and second convolution layer revealed recognizable motifs including strong TIS sequences (e.g. ACCAUG), stop codons (TAA, TGA, TAG), uORFs, and sequences composed of multiples of CG or AU likely involved in secondary structure formation (**Fig. 2F, Fig. S8 - S10**). Intriguingly, several filters did not fall into either of these categories and also did not match previously described PWMs for RNA binding proteins (Tomtom (*27*) and the Homo *sapiens* RBP database (*28*)), suggesting the possibility for previously undescribed regulatory interactions.

We then applied our method to transcripts bearing either pseudouridine (Ψ) or 1-methyl- pseudouridine (m^1^Ψ) instead of uridine (U) (**Fig. 2G**). These RNA modifications are widely used for mRNA therapeutics because they can increase mRNA stability and help modulate the host immune response (*29, 30*). We found that the model trained on the unmodified (U) library could explain 68% to 76% of the measured variability in the Ψ and m^1^Ψ polysome profiling data, respectively (**Fig. 2H**). Prediction accuracy could be further improved by training the models directly on data from the modified RNAs (the same held-out library sequences were used in all test sets to ensure consistency). This is likely due to the model learning the impact of Ψ and m^1^Ψ on the formation of secondary structure (31). Concordantly, mean ribosome load is more positively correlated with a UTR’s predicted minimum free energy (MFE) for Ψ (r = 0.56) and m^1^Ψ (r = 0.58) than for U (r = 0.43) (**Fig. 2I**).

As a further test of our model’s capabilities, we asked whether it could be used to engineer completely novel, functional 5′ UTRs. A tool capable of designing 5′ UTRs for a targeted level of protein expression would be a valuable asset for mRNA therapeutics and metabolic engineering. While there has been some success in this effort in prokaryotes and yeast (*32–34*), rational design of 5′ UTRs in human cells has not been demonstrated. We developed a genetic algorithm that iteratively edits an initial random 50-mer until it is predicted by the model to load a target number of ribosomes and thus achieve an intended level of translation activity (**Fig. 3A**) The model used for this process was developed before the model in Figure 2 and differs slightly in terms of network architecture (Online Methods) and performance (r^2^: 0.92) (**Fig. S11**) (**Online Methods**). We designed two sets of UTRs for testing. The sequences in the first set were designed to target MRLs of 3, 4, 5, 6, 7, 8, 9, and a no-limit maximum (**Fig. 3B**). The second set was designed to follow the step-wise evolution of a UTR. We set the algorithm to first select for sequences with low ribosome loading and then, after 800 iterations, to select for high ribosome loading. Each unique sequence generated by the algorithm as the UTR evolved was synthesized and tested (**Fig. 3C and Fig. S12**). We did this for 20 sequences where upstream AUGs were allowed and another 20 in which AUGs were not allowed.

**Figure 3.**
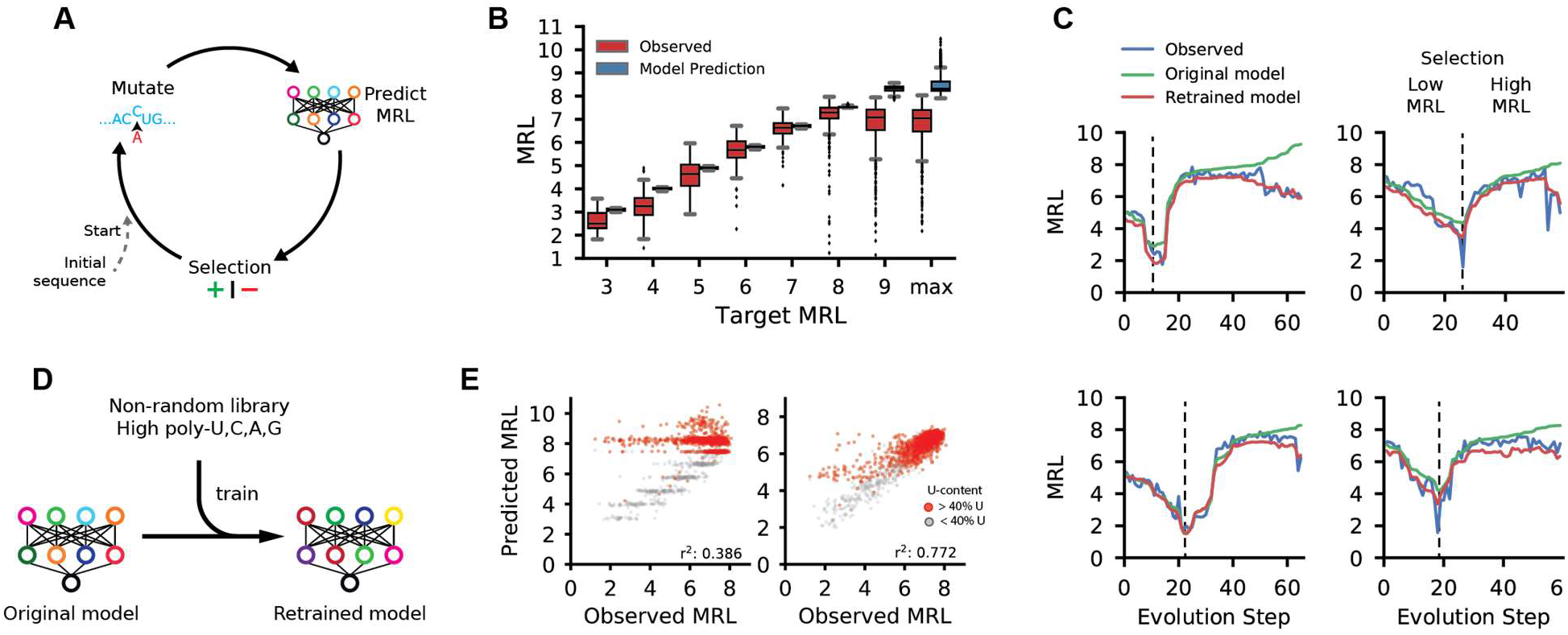
Design of new 5′ UTRs. **(A)** Diagram of a genetic algorithm that was used in conjunction with the 5′ UTR model to evolve sequences to target specific levels of ribosome loading. **(B)** Comparison between the predicted MRLs and observed MRLs for ~12,000 evolved 5′ UTRs for targeted ribosome loading. **(C)** Step-wise evolution analysis. Randomly initialized UTRs were first evolved for low ribosome loading and then for high ribosome (selection change at dashed line). Four out of 80 (**Fig. S10**). examples are shown. Examples on the left were permitted to have uAUGs while those on the right were not. Each unique sequence that was generated during the evolution process was synthesized and tested by polysome profiling. The original model prediction (green) and the observed MRL eventually diverge, but the predictions from the retrained model (red) more accurately reflect the data. **(D)** The original model is retrained using sequences from the designed library with high poly-U, C, A, and G stretches which occur rarely in the random library. **(E)** The accuracy of the retrained model increased significantly when predicting the high poly-U sequences (red) generated by the genetic algorithm (r^2^: 0.386 to 0.772).

Of the 12,000 total UTRs evolved for targeted expression in the first set, the median MRL for targets 3 through 8 followed the expected trend from low to high with low variability within each group. For the step-wise evolved UTRs in the second set, predicted MRLs (green) closely matched the trend of the observed (blue) along the trajectory. While we created sequences with high ribosome loading (**Fig. S13**), in both sets the prediction from the model and the observed MRL eventually diverged as the model produced UTRs with very high predicted MRLs. We suspected that the divergence between predictions and measurements at very high MRL values might reflect the unusual sequence composition of the maximally evolved UTRs which often contained multiple long stretches of poly-U – sequences rarely seen in the random library. We corrected the model by training it (**Fig. 3D**) for four additional iterations with 6,082 UTRs from the target MRL sub-library, which had a much higher frequency of homopolymers, and 2,695 previously unseen random UTRs. Reevaluation of held-out sequences from the ‘target MRL’ library showed a dramatic improvement in comparison to the original model (r^2^ from 0.386 to 0.772) (**Fig. 3E and Fig. S14**) as did the predicted loading of the step-wise evolved sequences (**Fig. 3C red line and Fig. S11**). Using this expanded dataset, we retrained the model in Figure 2, which showed increased accuracy with all sub-libraries and unchanged performance with random library sequences (**Fig. S15**). Due to this significant improvement, we used the retrained version of the model from this point on.

Can a model trained only on synthetic sequences predict the translation of human mRNAs from their 5′ UTR sequence? Assessing model performance on endogenous transcripts is challenging due to confounding contributions of 3′ UTRs and coding sequence lengths. As an alternative approach, we synthesized and tested via polysome profiling a 5′ UTR library consisting of the first 50 nucleotides preceding the start codon of 35,212 common human transcripts as well as 5′ UTR fragments carrying 3,577 variant sequences from the ClinVar database (35) that occur within these regions; the same eGFP context as the randomized library was used. Using the retrained model, we were able to explain 81% of the observed variation in MRL with the common and SNV 5′ UTR sequences (**Fig. 4A**) showing that, despite training on random sequences, the model was able to learn the cis-regulatory rules of human 5′ UTR sequences that lay directly upstream of a coding sequence.

**Figure 4.**
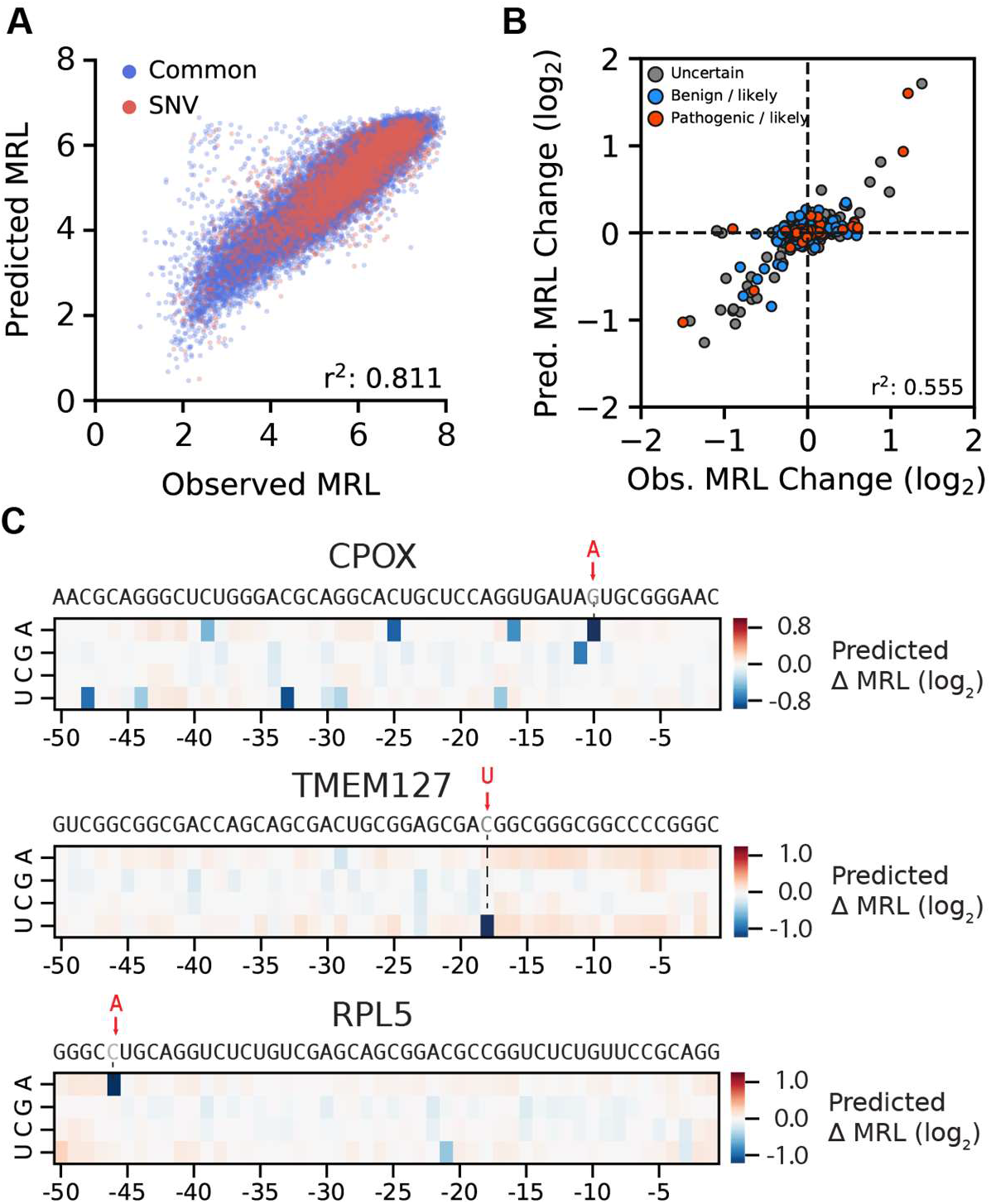
Model performance with human 5′ UTRs and SNVs. The first 50 nucleotides preceding the CDS of 35,212 human transcripts and an additional 3,577 UTRs with SNVs (ClinVar) were evaluated using our polysome profiling method with eGFP used as the CDS. **(A)** The retrained model could explain 81.1% of the observed variation in MRL. **(B)** The log_2_ change in MRL between an SNV and its common sequence was compared to the predicted change between the two (r^2^: 0.555). SNV classification labels are from the ClinVar database. **(C)** *In silico* saturation mutagenesis and model prediction of MRL change for all 5’ UTR variants of CPOX, TMEM127 and RPL5. The three annotated Clinvar variants, rs867711777 (CPOX, G > A), rs121908813 (TMEM127, C > U), and rs376208311 (RPL5, C > A), are predicted to have the most dramatic effect on ribosome loading.

Genetic variants play a major role in phenotypic differences between individuals (*36*) and how these sequences affect translation is only beginning to be understood (*37, 38*). But existing approaches to this problem, such as quantitative trait locus (QTL) analysis and genome wide association studies (GWAS) are limited to common variants and cannot scale to the enormous number of rare 5’UTR variants occurring in the human population. In contrast, a model-based approach can in principle be used to score the impact of any 5’UTR variant on translation. With this in mind, we investigated the model’s ability to predict the effect of disease relevant-variants by testing its performance in predicting the ribosome load change between pairs of wild-type (‘common’) and SNV-containing 5′ UTR sequences, measured as log2 difference. The majority of SNVs had little to no effect, but 47 had log2 differences greater than 0.5 or less than -0.5 (**Supplementary Table 1**). Overall, our retrained model could explain 55% of the observed MRL change (**Fig. 4B**) and accurately predicted the direction of change for 64% of the variants. Importantly, the model can explain 76% of the change of variants with log2 differences greater than 0.5 or less than -0.5 (**Fig. S16A**). As an example, one of the ClinVar variants with sizeable differences in MRL, rs867711777, is found in the 5′ UTR of the CPOX gene and shows a log2 difference of -0.89. The depletion of CPOX reduces heme biosynthesis and is the cause of hereditary coproporphyria (HCP) (*39*). The large MRL difference suggests that this SNV, labeled as uncertain in the ClinVar database, could be pathogenic. The variant rs376208311 lies in the 5′ UTR of the ribosomal subunit gene RPL5 and shows a -0.87 log2 difference in MRL. This variant is associated with Diamond-Blackfan anemia (BDA). One cause of the disease is a result of either the disruption or downregulation of RPL5 (*40*). Another SNV, rs121908813, is implicated in familial pheochromocytoma, a condition characterized by tumors found in the neuroendocrine system that secrete high levels of catecholamines (*41*). In our assay, the variant UTR shows a -1.5 log2 difference in MRL compared to the wild type 5′ UTR sequence. TMEM127 acts as a tumor suppressor and decreased expression of it could explain the observed pathogenicity of this variant. For the three examples, the model predicts that, of all possible variants, these specific SNVs, all of which introduce an upstream start codon, would most dramatically affect ribosome loading (**Fig. 4C**).

The method developed here, which combines polysome profiling of a randomized 5′ UTR library with deep learning, has provided a wealth of information detailing the relationship between the 5′ UTR sequence preceding a CDS and regulation of translation. The data and model enabled the quantitative assessment of secondary structure, uAUGs and uORFs, Kozak sequences, and other cis-regulatory sequence elements in the context of unmodified mRNA, Ψ, and m^1^Ψ-modified mRNA. The CNN trained on the data performed exceedingly well, explaining up to 93% of mean ribosome load variation in the test set and up to 81% of variation for 38,789 truncated human UTRs. The model also proved capable of predicting the effect of disease-relevant 5′ UTR variants on translation, even suggesting mechanisms of action. Importantly, predictions are not limited to common variants or even those that have been previously described; instead the model can be used to screen every possible SNV, insertion or deletion in the 50 bases upstream of the start codon – there are millions in the human genome - and select those for further study that have the strongest impact on ribosome loading and thus the highest likelihood of being pathogenic. Finally, using the model and a genetic algorithm, we were also able to engineer new 5′ UTR sequences for targeted ribosome loading, enabling even more forward-looking applications in synthetic biology and precision medicine.

## Acknowledgments

We would like to thank Alexander B. Rosenberg and Johannes Linder for helpful discussions on data analysis and modeling. We would also like to thank Melissa Moore, Andrew Hsieh, and Yiting Lim for constructive comments on the manuscript.

## Author contributions

P.J.S designed and performed experiments, performed data analysis and modeling, and wrote the manuscript. B.W. designed and performed polysome profiling assays and wrote the manuscript. D.R. performed fluorescence validation experiments. V.P. and I.M. wrote the manuscript. D.R.M. helped design polysome profiling. G.S. designed experiments and wrote the manuscript.

## Data and materials availability

All code is available at [https://github.com/pjsample/human_5utr_modeling] and data is available by request.

## Supplementary Materials

Materials and Methods

Figures S1-S17

Tables S1-S2

References (*42–45*)

